# Cluster Formation and Phase Separation Driven by Mobile Myosin Motors in the Motility Assay

**DOI:** 10.1101/2024.11.30.626189

**Authors:** Brandon Slater, Alfredo Sciortino, Andreas R. Bausch, Taeyoon Kim

## Abstract

Interactions between F-actin and myosin are critically important for a wide range of biological processes, including cell migration, cytokinesis, and morphogenesis. The motility assay with myosin motors fixed on a surface has been utilized for understanding various phenomena emerging from the interactions between F-actin and myosin. For example, F-actin in the motility assay exhibited distinct collective behaviors when actin concentration was above a critical threshold. Recent studies have performed the myosin motility assay on a lipid bilayer, meaning that myosin motors anchored on the fluid-like membrane have mobility. Interestingly, mobile motors led to very different collective behaviors of F-actin compared to those induced by stationary motors. However, the dynamics and mechanism of the unique collective behaviors have remained elusive. In this study, we employed our cutting-edge computational model to simulate the motility assay with mobile myosin motors. We reproduced the formation of actin clusters observed in experiments and showed that F-actin within clusters exhibits strong polar ordering and leads to phase separation between myosin motors and F-actin. The cluster formation was highly dependent on the average length and concentration of F-actin. Our study provides insights into understanding the collective behaviors of F-actins that could emerge under more physiological conditions.

---

Cells employ mechanical forces for a wide variety of physiological functions, such as cell migration (1), cytokinesis (2), and morphogenesis (3). In animal cells, the actin cytoskeleton is mainly responsible for generating these forces (4). Specifically, molecular interactions between actin filaments (F-actin) and myosin II motors in the actin cytoskeleton generate a large fraction of mechanical forces (5). F-actin is a semi-flexible biopolymer with a double-helical structure self-assembled by interactions between actin monomers (G-actin). Individual myosin II motors comprise two heads and one tail, and affinity between tails results in the assembly of myosin II into a higher order structure called a thick filament or a minifilament (6). Myosin heads on thick filaments or minifilaments bind to F-actin and walk toward the barbed end of F-actin. The walking of the myosin heads along F-actin is often described by the cross-bridge cycle (7). During the cross-bridge cycle, the myosin heads transition through different states at rates determined by mechanochemical properties (8).

Due to the importance of interactions between myosins and F-actins for biological processes, various experimental techniques have been developed to understand these interactions. One of the experimental techniques is the motility assay developed in 1986 (9, 10). In the traditional motility assays, myosin heads without a tail, called heavy meromyosin (HMM), are attached to a glass substrate (11). F-actins are propelled as the myosin heads walk toward to the barbed end of F-actins. Since myosin heads are fixed on surface, F-actins exhibit gliding motions. While the motility assay is different from physiological conditions, it has helped studying the interactions between myosin motors and F-actins in a well-controlled environment (12-14). In addition, previous studies using the motility assay have shown that F-actins exhibit interesting collective behaviors and various patterns, such as bands, clusters, density waves, and swirls, if F-actin density is above a critical threshold (15-18). At such a high density, repulsive forces acting between F-actins due to volume-exclusion effects are substantial enough to reorient and align F-actins. It was also shown that longer and denser F-actins lead to distinct bundle formation via more significant alignment between F-actins after collisions (19). Some of these studies have identified patterns in a dynamic steady state where F-actins either continuously leave and enter structures or continuously glide within structures.

In traditional motility assay experiments, F-actins can cross-over each other after binary collisions because they are free to move in the z-direction (i.e., one F-actin can cross-over the top of the other). However, during these cross-over events, the orientation of F-actin ends can change to some extent due to repulsive forces. As a result, there is relatively weak alignment between F-actins after collisions. One study found the multiple collisions are necessary to result in polar (parallel) or nematic (parallel or anti-parallel) ordering between F-actins, which is commonly seen in active matter systems (17). Furthermore, several studies demonstrated that structure formation and the type of filament ordering strongly depend on microscopic collision patterns between F-actins (17, 20-22). In motility assay experiments, the strength of volume-exclusion effects can be varied to some degree by changing the concentration of crowding agents that make F-actins stay near the surface via entropic effects (23). To vary collision patterns in a different manner, recent motility assay experiments used a lipid bilayer membrane beneath myosin heads rather than the glass surface, which allows myosin heads to move during interactions with F-actins (24, 25). This results in weaker repulsive forces between colliding F-actins because myosin motors anchored on the fluid-like membrane cannot exert strong propelling forces to F-actins. At the microscopic level, they found that filaments tended to stop upon collisions and align nematically. This led to the formation of collective structures, such as clusters or vortices with polar alignment between F-actins.

To understand the behaviors of F-actins emerging in the motility assay, computational models have been employed (26). In particular, agent-based models, which can explicitly describe individual filaments and their collision events, have been used popularly for simulating the motility assay. For example, one study showed that a variation in the persistence length (i.e., bending stiffness) of filaments or the extent of volume-exclusion effects can result in distinct collective behaviors, including flocks, polar streams, buckling bands, and spirals (27). However, most of the previous computational models used implicit motors, which means that F-actins are propelled by the direct application of forces rather than directly considering the molecular interactions between myosins and F-actins. Recently, we used our agent-based model with explicit motors to show how the formation of flocks, bands, and rings depends on volume-exclusion effects as well as the concentration, average length, and persistence length of filaments (28). Furthermore, we demonstrated that collective behaviors emerging from explicit motors are different from those from implicit motors.

In this study, we employed our agent-based model with explicit motors to demonstrate and analyze the collective behaviors of F-actins propelled by mobile motors. We found that patterns formed by collisions between F-actins are highly dependent on the mobility of motors. Under conditions where immobile motors form bundles or swarms, mobile motors lead to the formation of distinct aggregating structures with the pointed ends of F-actins orientated towards the center. After reaching a steady state, these aggregates are neither disassembled nor change in size. Interestingly, we found phase separation between motors and F-actins maintained robustly despite continuous diffusion of both motors and F-actins. We verified part of computational results using experiments obtained from the motility assay with the lipid bilayer membrane. Our study sheds light on the collective behaviors of F-actins that could emerge under more physiological conditions.

## RESULTS

### Mobile motors induced the formation of distinct actin clusters

First, we probed how the mobility of motors affects the pattern formation of F-actins. We performed simulations with different diffusion coefficients of motors, *D*_M_. We used 5 different values of 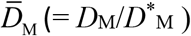 between 0 and 4, where *D*^M*^ is the reference value (Fig. 1). Higher 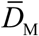 means more mobile motors, whereas 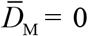 means stationary motors which are equivalent to those in the motility assay with a glass surface. We found that higher motor mobility led to the formation of distinct actin clusters (Fig. 1A, Movie S1). For instance, with 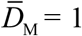 or 4, F-actins formed large, round structures, whereas, with 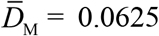 or 0, F-actins formed long, thin structures which we denote as bundles. These actin clusters are reminiscent of aggregated structures observed in our motility assay experiments performed with a lipid bilayer (24), meaning that our model is able to capture the collective behaviors of F-actins driven by mobile motors observed in experiments. We also quantified the speed of F-actins and motors as a function of 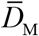As expected, higher motor mobility led to higher motor speed but lower F-actin speed (Fig. 1C). Motors experience reaction forces when they apply propelling forces to F-actins. Thus, as motors are more mobile, they are pushed back more by F-actins, and thus F-actin speed is reduced.

**Figure 1.**
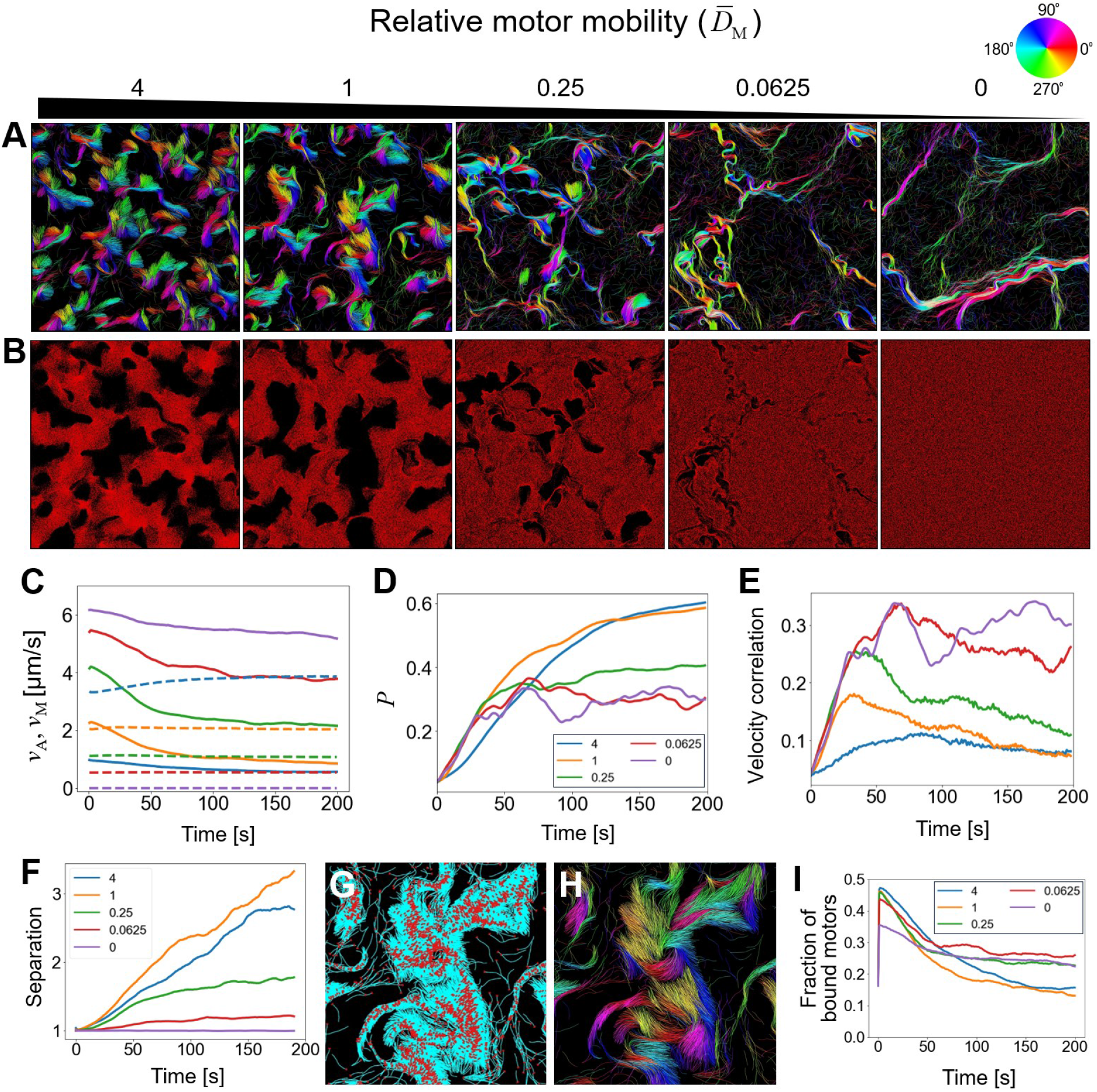
Mobile motors led to the formation of distinct actin clusters. (A, B) Snapshots showing (A) actin structures and the orientations of F-actins and (B) the spatial distribution of m<u>o</u>tors at 200 s with different motor mobility defined by the relative diffusion coefficient of motors 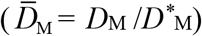. With larger motor mobility, separated cluster structures tended to form more likely, and phase separation between F-actins and motors was more apparent. (C) Average speed of F-actins (*v*_A_, solid lines) and motors (*v*_M_, dashed lines). With higher motor mobility, *v*_M_ was higher, whereas *v*_A_ was smaller. (D) Polar order parameter measured using nearby F-actins. Higher motor mobility led to more nematic ordering. (E) Velocity correlation between neighboring F-actins. The velocity corre<u>l</u>ation tended to be lower with higher motor mobility. The legend in (C) indicates the values of 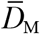, and it is shared with (B) and (D). (F) Extent of phase separation between F-actins and motors. <u>C</u>ases with more mobile motors showed greater phase separation. (G) Clusters in the case with 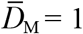. The pointed ends of F-actins visualized in red, whereas the other parts of F-actins are shown in cyan. (H) There are the +1/2 local nematic defects in cluster structures. (I) Fraction of motors bound to F-actins.

### F-actins in the clusters underwent the polar ordering but showed poor velocity correlations

We quantified the polar and nematic order parameters between neighboring F-actins pairs. The polar order parameter (*P*) indicates the extent of parallel alignment, whereas the nematic order parameter (*S*) represents the degree of both parallel and anti-parallel alignments. For example, both *S* and *P* are close to 1 for a system with parallelly aligned F-actins, whereas only *S* is close to 1 for a system with the mixture of parallelly and anti-parallelly aligned F-actins. *S* was high in all cases regardless of motor mobility (Fig. S2A). However, *P* showed dependence on motor mobility. When motor mobility was high (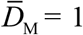 or 4), *P* was high, meaning that most of F-actins were parallelly aligned in the clustering structures. (Fig. 1D). By contrast, when motor mobility was low (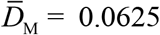 or 0), *P* was lower, implying the existence of both parallel and anti-parallel alignments in the bundle structures. This indicates that higher motor mobility specifically leads to the polar ordering of F-actins.

As another indicator for F-actin alignment, we analyzed a velocity correlation between neighboring F-actin pairs (Fig. 1E). It was found that increased motor mobility led to a lower velocity correlation. Interestingly, the velocity correlation increased rapidly at the beginning of the simulations and then gradually decreased later in all cases with mobile motors. When the motors were immobile, this tendency was not observed at all. These observations imply that bundle formation results in a higher velocity correlation between neighboring F-actins; F-actins forming bundles keep interacting with motors and consistently gliding in directions relatively parallel to the axis of the bundles (Movie S2, right). In cases with actin clusters, F-actins were aligned mostly in a parallel manner at the beginning of cluster formation, but they showed negligible movement after cluster formation (Movie S2, left). It means that the velocity vector of F-actin may not be parallel to the current orientation vector of F-actin. Thus, the velocity correlation between neighboring F-actins in the clusters decreased over time although they were still parallel to each other.

Additionally, we analyzed the quantity of filaments entering or leaving either clusters or bundles to understand their difference. First, we selected three representative clustering regions (Fig. S3A) and three representative bundle regions (Fig. S4A). In the case of clusters, while the noise was quite high, both the number of filaments entering and leaving the clustering regions decreased over time and approached zero (Figs. S3B-G). The net number of F-actins within the clustering regions tended to increase over time. This indicates that clusters act as a trap or a sink for F-actins. In the case of bundles, the number of F-actins entering and leaving bundling regions remained high or increased over time (Figs. S4B-G). However, the net number of F-actins within the bundling regions showed fluctuations or large drops. This implies that F-actins within bundles showed continuous movements rather than staying in the same regions. These observations imply a fundamental difference between clusters and bundles.

### Polar ordering of F-actins leads to phase separation between motors and F-actins

We further investigated why F-actins showed negligible motions after cluster formation. It was found that the formation of actin clusters led to phase separation between F-actins and motors. The extent of phase separation was the highest in the case with 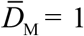 that showed the largest cluster formation and the highest *P* (Figs. 1A, B, D, F). In the case with 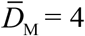showing slightly lower *P* and smaller F-actin clusters, less phase separation was observed than that in the case with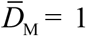. By contrast, with low mobility (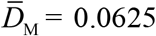or 0), little to no phase separation was observed between F-actins and motors. We delved into the molecular origin of the phase separation emerging with mobile motors. We found that the pointed ends of F-actins were located mostly near the center of clusters (Figs. 1G and S2B). Similar to our previous observation (24), we observed the formation of “+1/2 local nematic” defects (Fig. 1H). These configurations resulted in the polar alignment of F-actins, and the distinct F-actin orientation led to phase separation between F-actins and motors. If motors diffuse toward and subsequently bind to F-actins in a cluster, they will walk towards the barbed ends of F-actins rather than propelling F-actins because the pointed ends are surrounded by many other F-actins belonging to the same cluster. As a result, motors will be pushed out of the clusters. This leads to the stable phase separation observed in our simulations.

To further quantify the phase separation between F-actins and motors, we also analyzed the fraction of motors bound to F-actins (Fig. 1I). The fraction spiked at initial times and then decreased substantially to a steady level. With higher motor mobility, the steady level was lower, meaning that there were fewer motors interacting with F-actins due to the phase separation between motors and F-actins.

### The length and concentration of F-actins affected the collective behaviors of F-actins

In our previous computational study (28), we have shown the significant effects of a change in the average length or concentration of F-actins on the collective behaviors of F-actins driven by stationary motors. It is expected that these parameters would also influence the collective behaviors of F-actins propelled by mobile motors. First, we varied the average F-actin length, <*L*_f_>, from its reference value, 2.58 µm. <*L*_f_> was adjusted by varying the actin nucleation rate during the initiation phase. When <*L*_f_> was lower, the formation of separated actin clusters was very obvious (Fig. 2A). The cluster formation was also supported by high *P*, significant separation between F-actins and motors, the low fraction of F-actin bound motors, and the velocity correlation showing an initial increase followed by a gradual decrease (Figs. 2B-F). By contrast, when <*L*_f_> was higher, F-actins did undergo cluster formation. It is generally harder for longer F-actins to be reoriented, so they instead tended to form closed loop structures (i.e., rings). Note that cluster formation requires the polar ordering of relatively straight F-actins. Cases with high <*L*_f_> showed similarities to those with low motor mobility which showed bundle formation, in terms of low *P*, significantly less separation between F-actins and motors, the larger fraction of F-actin-bound motors, and the velocity correlation that initially increased and then reached a steady state.

**Figure 2.**
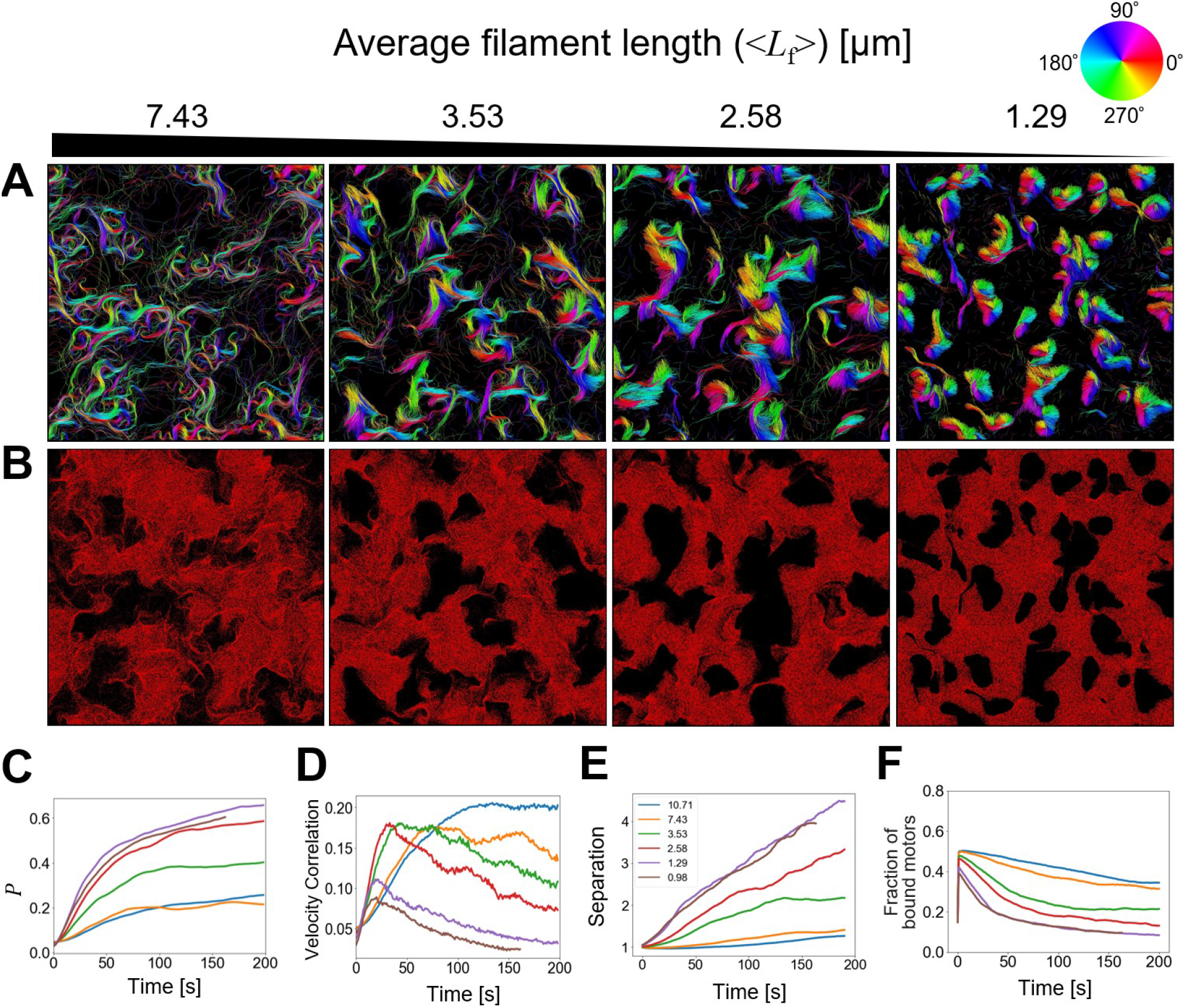
Distinct actin clusters were more likely to form with shorter F-actins. (A, B) Snapshots at 200 s representing (A) actin structures and F- actin orientations and (B) the spatial distribution of motors for different average F-actin lengths (<*L*_f_>). With shorter F-actins, clusters were separated more, and phase separation between F-actins and motors was greater. (C) Polar order parameter between nearby F-actins, (D) velocity correlation between neighboring F-actins, (E) phase separation between F-actins and motors, and (F) fraction of motors bound to F-actins, with different <*L*_f_>. With shorter F-actins, nematic ordering of filaments was more significant, but the velocity correlation was lower. In addition, phase separation between F-actins and motors was more severe. The legend in (E) indicates the values of <*L*_f_>, and it is shared with (C), (D), and (F).

We also varied actin density defined by the number of actin monomers per µm^2^, whereas motor density was fixed in all simulations. When the actin density was reduced to a quarter of the reference case, few collective behaviors were observed (Fig. 3A). The collective behaviors of F-actins are driven by collisions between F-actins. With very low actin density, average distance between F-actins is too long to induce collisions between F-actins frequently enough to result in any collective behavior. Due to lack of cluster formation, we observed low *P*, negligible F-actin-motor separation, and the large fraction of F-actin-bound motors (Figs. 3B-F). With higher actin density than the reference case, the formation of separated actin clusters was more pronounced because there were more opportunities for F-actins to collide to each other. These cases showed high *P*, significant separation between F-actins and motors, and the small fraction of F-actin-bound motors.

**Figure 3.**
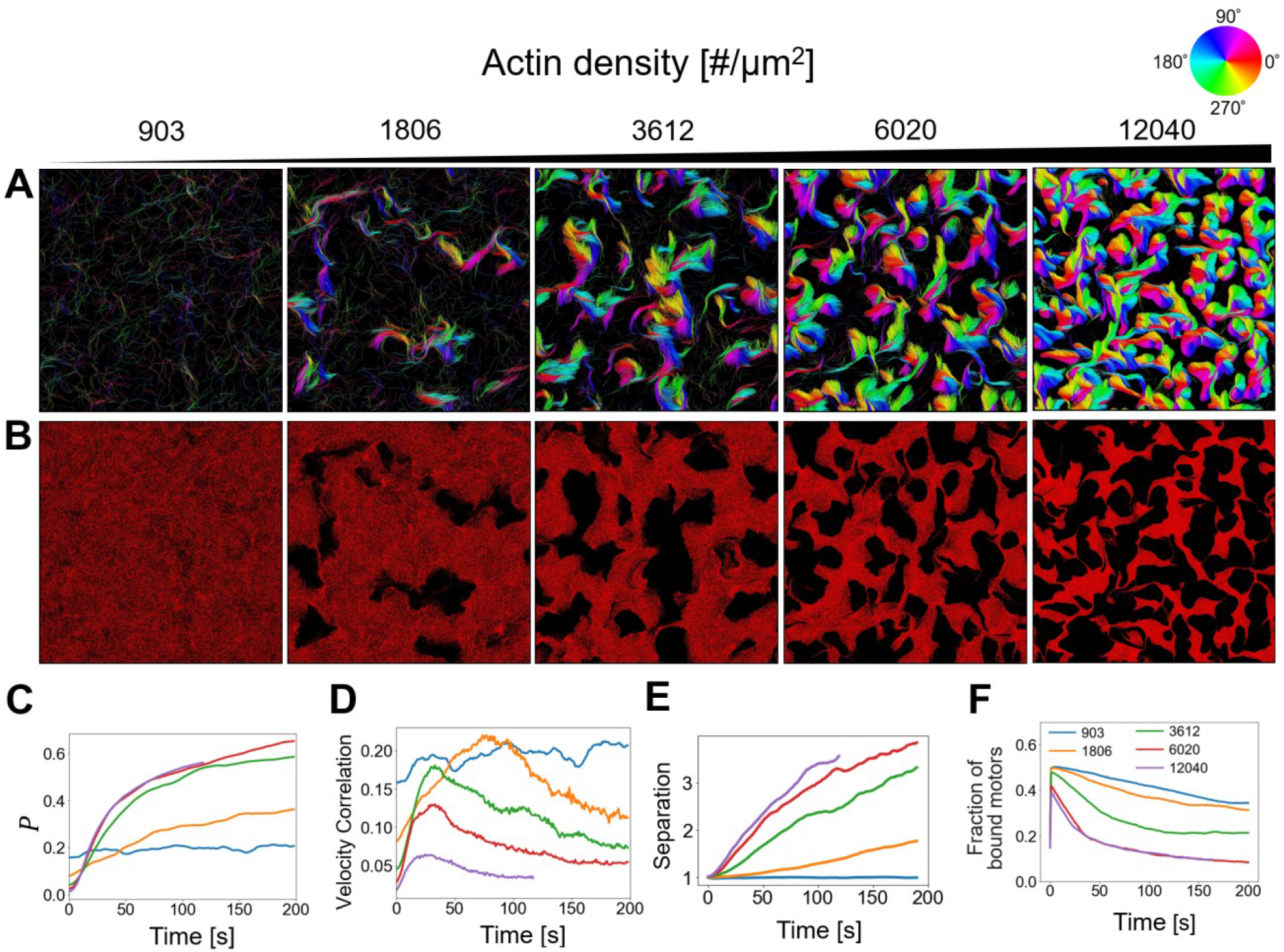
Higher actin density mediates more distinct cluster formation and greater phase separation. (A, B) Snapshots at 200 s showing (A) actin structures with visualization of filament orientations via color scaling and (B) the spatial distribution of motors, with varied actin density defined by the number of actin monomers per µm^2^. Phase separation and cluster formation were more severe with higher actin density. (C) Polar order parameter between neighboring F-actins, (D) velocity correlation between nearby F-actins, (E) phase separation between F-actins and motors, and (F) fraction of motors bound to F-actins, with five different values of actin density shown in the legend of (E). With larger actin density, nematic ordering of F-actins was more significant, whereas the velocity correlation was smaller. In addition, phase separation between F-actins and motors was greater. The legend in (E) indicates the values of actin density, and it is shared with (C), (D), and (F).

**Figure 4.**
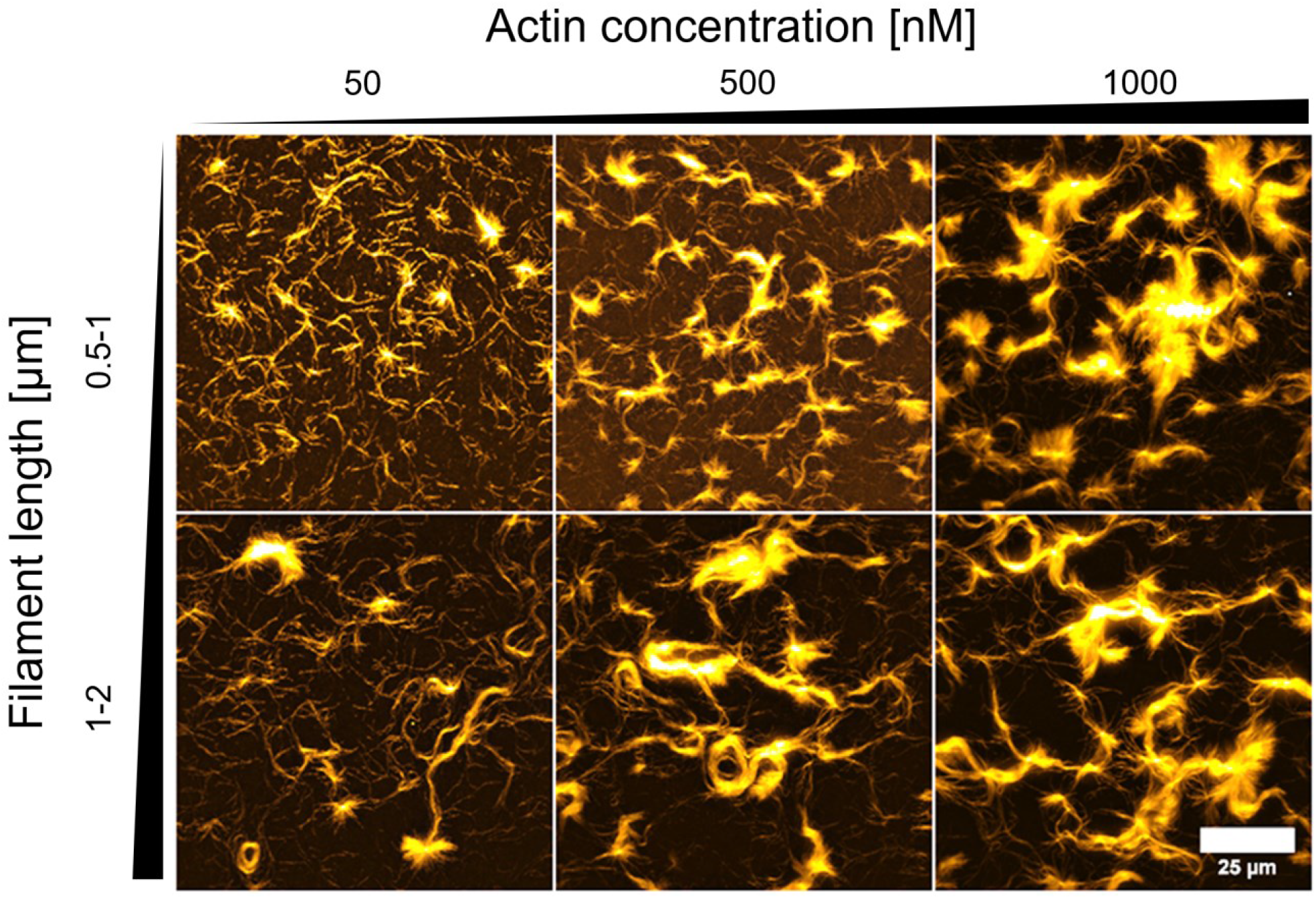
Experimental observations verify simulation results. The experiments were performed using the motility assay with mobile myosin motors bound to a lipid bilayer, with 3 different actin concentrations and 2 different filament lengths. Overall, higher actin concentration and shorter filament length resulted in more distinct cluster formation, which is consistent with simulation results.

### Experimental results support our findings

To verify our simulations, we quantitatively compared them to experimental results obtained with mobile motors. The experiments were performed as in our previous work (24) by binding biotinylated HMM motors to a supported lipid bilayer and then flowing in stabilized F-actins at different concentrations. The average F-actin length is controlled during polymerization via different concentrations of the actin-severing protein, gelsolin. As in simulations, motor concentration was kept constant, whereas the concentration and length of F-actins were varied in experiments.

The experiments clearly showed the presences of both elongated structures (bundles) and high-density clusters. With lower actin concentration, shorter F-actins (∼0.5-1 µm) showed a propensity to form clusters, whereas longer F-actins (∼1-2 µm) formed elongated bundles. With higher actin concentration, cases formed clusters regardless of average F-actin length. While we were not able to observe phase separation between F-actins and motors in this experimental setup, the phase separation has been reported recently in the microtubule-based motility assays (21). These results qualitatively match the simulation results. Overall, these provide verifications for the simulation results.

## DISCUSSION

It has been well-established that active matter systems exhibit collective behaviors when the concentration is above a critical threshold. For instance, in the myosin motility assay, it has been shown that F-actins exhibit distinct collective behaviors at high actin concentration. However, recent studies have shown that biopolymers propelled by mobile motors show distinctly different patterns than those with fixed motors. For example, our previous experimental study performed the motility assay with myosin motors bound to a lipid bilayer instead of a glass surface (24). Other studies have performed the motility assay using microtubules with mobile kinesin motors attached to a lipid bilayer (21, 34). If motors are fixed in place, F-actins are more likely to cross-over each other after collisions due to strong propelling forces. However, mobile motors lead to a regime where steric hindrance between F-actins significantly affects alignment. For example, mobile motors result in the formation of clustering patterns that show polar ordering (strong parallel alignment). Interestingly, this occurred in a regime where nematic alignment (parallel and anti- parallel alignment) occurs when motors are fixed.

In this study, we employed our agent-based computational model to show how mobile motors lead to the formation of distinct clusters. The key advantage of our model is the explicit representation of myosin motors. In most of the previous models, myosin motors were represented implicitly by directly applying a constant propelling force to filaments. While this reduces the computational cost, it also reduces the physiological relevance and may lead to artifacts. For instance, in our previous study, we found that explicit motors lead to more rigorous results, which cannot be reproduced with implicit motors (28).

First, we varied the mobility of the myosin motors. To do this, we changed the strength of the drag coefficient of the myosin motors with the surrounding environment. A larger drag coefficient leads to less more motor mobility (and vice-versa). We found that more mobile motors lead to the formation of distinct clusters. When motor mobility is very low, we observed F-actins formed bundles. Additionally, we found that F-actins within clusters show strong polar ordering while F-actins within bundles show nematic ordering. This is in agreement with recent experimental studies (24). Interestingly, F-actins within clusters show poor velocity correlations.

Additionally, our findings suggest that F-actins clusters act as trap where filaments tend to enter clusters and are unable to leave. Within clusters, F-actins are orientated with the pointed-end towards the center. As a result, any myosin motors within cluster are pushed outwards. This leads to a phase separation between motors and filaments. Finally, we showed how the average length and concentration of F-actins affect the collective. We found that shorter filaments would more easily undergo polar alignment to form clusters. Additionally, when F-actin concentration was higher, F-actins more readily formed clusters. Our computational results were verified by qualitatively comparing with experiments.

While the motility assay has been widely used to study the interactions between myosin motors and F-actins, it can lack physiological context. By incorporating the mobile motors, we may improve the physiological relevance. Furthermore, there are several limitations of our model. First, we didn’t include any repulsive force between motors. Since the motor density is very higher, the phase separation within clusters may be reduced as myosin motors repel each other. Additionally, our model did not account of serving, which some experimental studies have established effect collective behavior.

## CONCLUSIONS

Our study provides insights into how mobile motors lead to the formation of distinct actin clusters. Using a rigorous agent-based model, we showed how F-actins undergo polar alignment to clusters and show clusters are stabilized by phase separation between F-actins and motors. Additionally, we showed how two parameters, average filament length and filament concentration, affect the collective patterns formed.

## METHODS

### Model overview

Our agent-based computational model for modeling the motility assay is based on Brownian dynamics. The model is well-established and has been used in our previous studies (29- 31). Details about the model and parameter values used in this study are explained in Materials and Methods and Supporting Information. In the model, F-actins and motors are simplified via cylindrical segments (Fig. S1A). At each timestep, the displacements of F-actins and myosin motors in x and y directions are determined using the Langevin equation and the Euler integration scheme, whereas their z positions are unchanged. Extensional, bending, and repulsive forces govern the mechanical behaviors of F-actins, whereas motors are regulated by extensional forces only. The mobility of motors is varied by imposing a different value for the drag coefficient of motors; a higher drag coefficient corresponds to less mobility. During the initiation phase, actin undergoes nucleation and polymerization to form F-actins uniformly along xy plane at z 13.5 nm in a three-dimensional rectangular domain (5 ×5 × .1 µm) with the periodic boundary condition in the x and y directions. A large number of motors are also randomly positioned along xy plane at z. Motors can bind to F-actins but do not walk along F-actins during the initiation phase. After the initiation phase, motors start walking toward the barbed end of F-actins, leading to the movements of F-actins (and motors in cases with mobile motors).

### Implementation of motors in the model

In typical motility assay experiments, the glass surface is fully covered with myosin heads. Thus, the areal density of myosin heads is extremely high, which explains why most of the previous computational models for the motility assay used implicit motors. Since implicit motors propel F-actins directly using a constant force, the computational cost is significantly less. Our model employs explicit motors to rigorously account for molecular interactions between myosin motors and F-actins. However, some extent of simplification is inevitable. In our model, each motor is represented by a single segment that represents *N*_H_ myosin heads in the motility assay experiments, in terms of a stall force and kinetics. The most important kinetic parameters of motors are the rates of binding, unbinding, and walking. It is assumed that motors arms can bind to specific binding sites located on F-actins every 7 nm at a rate of *k*_+,M_ 4 *N*_H_ s^-1^. We assumed that *N*_H_ 4, which is moderate simplification of 4 myosin heads into one motor segment. The walking (*k*_w,M_) and unbinding (*k*_u,M_) rates are determined by the parallel cluster model (PCM) as in our previous studies (32, 33). The PCM assumes that myosin heads can exist in three different mechanochemical states, and transitions between the states are described by 5 transition rates (Table S2). The PCM can predict the unbinding and walking rates of a small ensemble with several myosin heads. Further details about the benchmarking and verification of the PCM can be found in our previous study (29).

To mimic mobile motors anchored on the fluid-like lipid bilayer used in our recent motility assay experiments (24), motors in our model are allowed to move against a drag force which is directly proportional to the drag coefficient (*ζ*_M_) and the velocity of motors. To change motor mobility, we vary *ζ*_M_. When *ζ*_M_ is higher, motors move less in response to the same applied force. In some of the simulations, motors are fixed in space to serve as control cases. In those cases, *ζ*_M_ is equivalent to an infinitely large value, ∞. To use a more intuitive value, we calculate the diffusion coefficient of motors using *ζ*_M_, *D*_M_ *k*_B_*T* / *ζ*_M_, where *k*_B_*T* is thermal energy. The reference value of *D*_M_ is 5.11×1 ^-14^ m^2^/s. To verify a variation in motor mobility, simulations were run without F-actins, and then the mean squared displacement of motors was quantified (Fig. S1B).

### Simulation setup

During the initiation phase, actin undergoes nucleation and polymerization to form F-actins in the absence of depolymerization within a thin three-dimensional computational domain (5 ×5 × .1 µm). The z-position of F-actins is fixed at z 13.5 nm. Motors are randomly positioned in the x and y directions at z µm. Note that the distance of 13.5 nm between the xy plane where F-actins are located and the xy plane where motors exist corresponds to the equilibrium length of motor segments. The periodic boundary condition is applied in the x and y directions. Motors bind to F-actins but do not walk along F-actins during the initiation phase. After the initiation phase, all actin dynamics stops, preserving the same distribution of filament length till the end of the simulation. After binding to F-actins, motors start walking toward the barbed end of F-actins, leading to the movements of F-actins (and motors in cases with mobile motors). The actin concentration (*C*_A_) was varied between 15 µM to 2 µM, which is equivalent to actin density between 9 3 to 12, 4 in the unit of the number of actin monomers per µm^2^. By contrast, the motor concentration (*C*_M_) was fixed at 6 µM in all simulations, which corresponds to 451,648 motors in the entire domain or ∼181 motors per µm^2^. Since each motor represents 4 myosin heads, effective myosin concentration is 24 µM (or equivalently 724 myosin per µm^2^), and the effective number of myosin heads is ∼1.8 million.

### Analysis of order parameters

To quantify the collective behavior of F-actins, we calculate “nematic” and “polar” order parameters by considering all F-actin pairs located within 1 µm. The difference between the two order parameters is whether parallel alignment of F-actins is distinguished from anti-parallel alignment. The nematic order parameter indicates the extent of F-actin alignment (regardless of whether the alignment is parallel or anti-parallel), and it can be computed, using the following equation for the two-dimensional space:

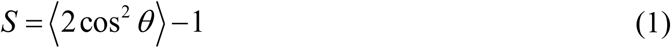

where *θ* is an angle between º and 18 º measured between pairs of neighboring F-actins. When *S* = 0, the ordering of F-actins is isotropic (i.e., no filament alignment). By contrast, when *S* 1, F-actins are ordered with perfect parallel/anti-parallel alignment. If most of the F-actins are orientated orthogonally, it is possible that *S* becomes negative. The polar order parameter is indicative of the extent of only parallel alignment, and it is calculated as follows:

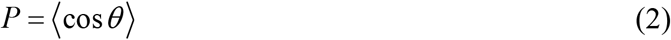

*P* ranges between -1 and 1. *P* and *P* 1 indicate no filament alignment and perfect parallel alignment, respectively.

### Quantification of separation and interactions between F-actins and myosin motors

The degree of separation between F-actins and motors is quantified as follows. First, the computational domain is divided into *N*_g_× *N*_g_ grids in the x and y directions. Through trial-and-errors, we found that 1 is a reasonable value for *N*_g_. In each grid, the total numbers of motors and actin segments are calculated and multiplied together, and the average of the multiplied values over all grids, *M*, is calculated. Using the initial state with uniformly distributed motors and F-actins in each simulation, the initial value, *M*, is found. Then, *M* is divided by *M* as a measure of phase separation between F-actins and motors over time.

A velocity correlation between neighboring F-actins is also calculated. First, the velocity of each endpoint of actin segments is calculated every 1 s. As in our previous studies (28), a correlation between nearby endpoints located within the cut-off radius of 1 µm is calculated:

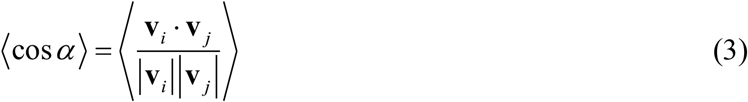

where *i* and *j* are the indices of actin segment endpoints, and α corresponds to an angle between two velocity vectors of the endpoints. The ensemble average is calculated using all pairs located within 1 µm. The velocity correlation close to 1 indicates that F-actins are moving in the same direction. The velocity correlation close to -1 represents that F-actins are moving in opposite directions. The velocity correlation close to indicates that the movements of F-actins are not significantly related to each other.

### Experiments

We performed the motility assay experiments using myosins anchored to a lipid bilayer membrane, with different sets of actin concentration and filament length. Detailed descriptions about experiments are included in Supporting Information.

## AUTHOR CONTRIBUTIONS

B.S. and T.K. designed the project. B.S. performed simulations and analyzed data obtained from the simulations. T.K. supervised all computational studies. A.S. conducted experiments and generated images from the experiments. A.R.B. supervised all experimental studies. All the authors participated in writing the manuscript.

## DATA AVAILABILITY STATEMENT

The data that support the findings of this study are available from the corresponding authors upon reasonable request.

## CONFLICT OF INTEREST DISCOSURE

The authors declare no conflict of interest.

