## Supporting Information for "Cluster Formation and Phase Separation Driven by Mobile Myosin Motors in the Motility Assay"

### SUPPORTING TEXT

#### Brownian dynamics via the Langevin equation

The velocities of the endpoints of actin segments at each time step are calculated by the Langevin equation without consideration of inertia:

$$\mathbf{F}_i - \zeta_i \frac{\mathrm{d}\mathbf{r}_i}{\mathrm{d}t} + \mathbf{F}_i^{\mathrm{T}} = 0 \tag{S1}$$

where the subscript $i$ represents $i$th endpoint, $\mathbf{F}_i$ is a deterministic force, $\zeta_i$ is a drag coefficient, $\mathbf{r}_i$ is position, $t$ is time, and $\mathbf{F}^{\mathrm{T}}_i$ is a stochastic force satisfying the fluctuation-dissipation theorem:

$$\left\langle \mathbf{F}_i^{\mathrm{T}}(t)\mathbf{F}_j^{\mathrm{T}}(t)\right\rangle = \frac{2k_{\mathbf{B}}T\zeta_i\delta_{ij}}{\Delta t}\boldsymbol{\delta} \tag{S2}$$

where $\delta_{ij}$ is the Kronecker delta, and $\boldsymbol{\delta}$ is a unit second-order tensor. Time step ($\Delta t$) is $1.15\times10^{-5}$ s. The approximated form of the drag coefficient for a cylindrical object is used:

$$\zeta_i = 3\pi\mu r_{\mathrm{c},i}\frac{3+2r_{0,i}/r_{\mathrm{c},i}}{5} \tag{S3}$$

where $\mu$ is the viscosity of the medium, and $r_{0,i}$ and $r_{\mathrm{c},i}$ are the length and diameter of a cylindrical segment, respectively. The positions of the endpoints of cylindrical segments are updated at each time step via the Euler integration scheme:

$$\mathbf{r}_i(t+\Delta t) = \mathbf{r}_i(t) + \frac{\mathrm{d}\mathbf{r}_i}{\mathrm{d}t}\Delta t = \mathbf{r}_i(t) + \frac{1}{\zeta_i}\left(\mathbf{F}_i + \mathbf{F}_i^{\mathrm{T}}\right)\Delta t \tag{S4}$$

#### Deterministic forces

Deterministic forces include extensional, bending, and repulsive forces. The extensional force acting on actin segments is determined by the following harmonic potential:

$$U_{\mathrm{s,A}} = \frac{1}{2}\kappa_{\mathrm{s,A}}(r - r_{0,\mathrm{A}})^2 \tag{S5}$$

where $\kappa_{\mathrm{s,A}}$ is extensional stiffness, and $r$ and $r_{0,\mathrm{A}}$ are the instantaneous and equilibrium length of actin segments, respectively. The extensional force acting on motor segments is also determined by a similar harmonic potential:

$$U_{\mathrm{s,M}} = \frac{1}{2}\kappa_{\mathrm{s,M}}(r - r_{0,\mathrm{M}})^2 \tag{S6}$$

where $\kappa_{\mathrm{s,M}}$ and $r_{0,\mathrm{M}}$ are extensional stiffness and equilibrium length, respectively.

The bending force acting on F-actin is determined by the following harmonic potential:

$$U_{\mathrm{b,A}} = \frac{1}{2}\kappa_{\mathrm{b,A}}\left(\theta - \theta_{0,\mathrm{A}}\right)^2 \tag{S7}$$

where $\kappa_{\mathrm{b,A}}$ is bending stiffness, and $\theta$ and $\theta_0$ are instantaneous and equilibrium angles formed by adjacent actin segments, respectively.

The repulsive force acting between two overlapping actin segments is governed by the following harmonic potential:

$$U_{\mathrm{r}} = \begin{cases} \frac{1}{2}\kappa_{\mathrm{r}}\left(r_{12} - r_{\mathrm{c}}\right)^2 & \text{if} \quad r_{12} < r_{\mathrm{c}} \\ 0 & \text{if} \quad r_{12} \geq r_{\mathrm{c}} \end{cases} \tag{S8}$$

where $\kappa_{\mathrm{r,A}}$ is repulsive strength, $r_{12}$ is a minimum distance between a pair of actin segments, and $r_{\mathrm{c}}$ is used as a cut-off distance above which the repulsive force becomes zero. $r_{\mathrm{c}}$ is equal to the diameter of actin segments, $r_{\mathrm{c,A}}$.

#### Experimental setup

##### *Proteins and buffers*

The main Assay Buffer (A-Buffer) is 25 mM Imidazole (pH 7.4), 4 mM $MgCl_2$, 25 mM KCl, and 1 mM EGTA. The actin polymerization buffer is F25 Buffer with 50 mM Tris (pH 7.5), 2 mM $MgCl_2$, 0.5 mM ATP, 0.2 mM $CaCl_2$, 25 mM KCl, and 1mM DTT. Actin and skeletal muscle myosin were purified from rabbit skeletal muscle. No rabbit was directly involved in the study. Monomeric actin was stored at 4 °C in G-Buffer (2 mM Tris, 0.2 mM ATP, 0.2 mM $CaCl_2$, 0.2 mM DTT, and 0.005% $NaN_3$ at pH 8.0). Heavy meromyosins (HMM) were prepared by dialyzing ground rabbit skeletal muscle against the Myosin-buffer (0.6 M KCl, 10 mM $KH_2PO_4$, and 2 mM DTT) at 4 °C (1). Gelsolin was purified from adult bovine serum (Sigma Aldrich). Alexa Fluor 488 phalloidin and streptavidin are purchased from Thermo Fisher. Lipids (DOPC, DSPE-PEG [2000] Biotin, Texas Red DHPE) are purchased from Avanti.

##### *Preparation of actin fragments and streptavidin-bound motors*

Actin is incubated in F25 buffer together with labelled phalloidin (1:2 phalloidin:actin) and with the desired concentration of gelsolin to vary average filament length. Three different batches of filaments are prepared: short (1:50 gelsolin:actin M/M), medium (1:190 gelsolin:actin M/M) and long (1:500 gelsolin:actin M/M). Filaments are polymerized at room temperature for 45 minutes and then stored on ice and used within a week.

HMMs are biotinylated, incubated with streptavidin in 1:1 M/M ratio, and then centrifuged together with a 5-fold amount (M/M) of pre-polymerized actin (45 min in F25 with phalloidin), at 4 °C at 350000xg for 25 minutes. The pellet is then discarded to remove enzymatically inactive head. The active motors are used within a day.

*Bilayer formation*

In short, a supported lipid bilayer (SLB) is formed with 98 % M/M DOPC, 2 % M/M DSPE-PEG [2000] Biotin, and 0.05 % M/M Texas Red DHPE on an observation chamber consisting of a coverslip and a glass slide separated by three layers of parafilm. Coverslips are hydrophilized using the Jelight UVO-Cleaner.

*Observations*

The observation chamber is then washed successively with A-Buffer, then 100 nM of streptavidin-functionalized HMM motors (incubated for 5 minutes), and then by actin filaments at desired concentration (incubated for 3 minutes). After actin incubation, the chamber is washed with A-Buffer, and snapshots are acquired to measure the surface density. The experiment is then started by washing with 2 mM ATP in A-Buffer together with 0.2 % Methylcellulose and an oxygen scavenging system (Glucose-oxidase from Sigma and catalase from Fluka). Finally, the chamber is sealed with vacuum grease and observed with a Total Internal reflection fluorescence (TIRF) microscope (Leica DMi8 with Infinity Scanner using a 100× TIRF objective). Snapshots are taken in different areas of the sample after approximately 30 minutes from ATP addition.

*Estimations*

The surface density (filaments/$\mu m^2$) is estimated by counting filaments in a 5×5 $\mu m^2$ region. The molar concentration is estimated using the mean length computed with Fiji's line-tool, divided by an average monomer size of value of 2.6 nm/monomers, which takes into account the helical pitch of F-actin.

#### SUPPORTING FIGURES

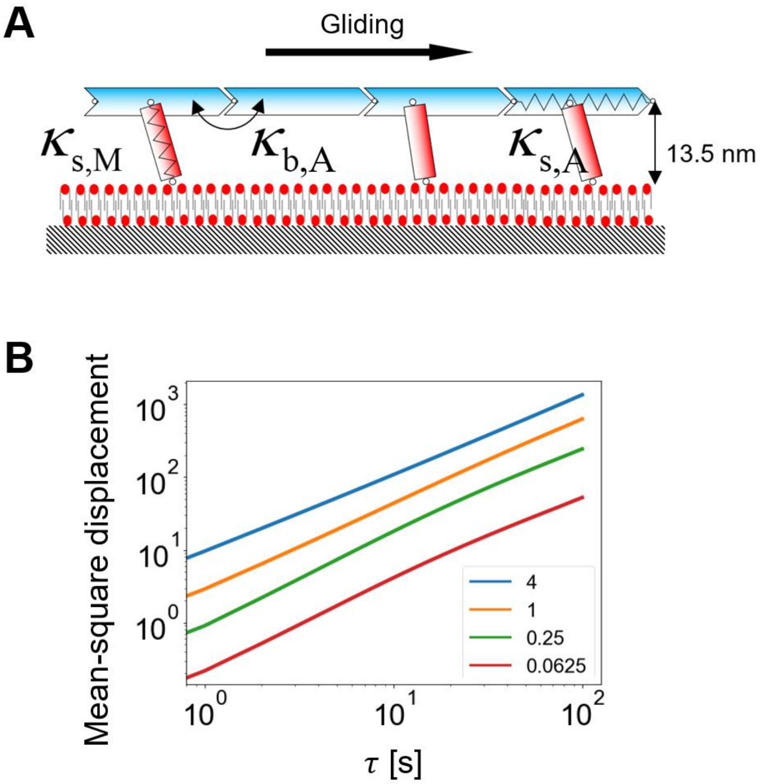


**Figure S1. Agent-based model.** (A) Schematic diagram showing our motility assay system where actin filaments are propelled by mobile motors. $\kappa_{s,A}$ and $\kappa_{b,A}$ are the extensional and bending stiffness of F-actin, respectively, and $\kappa_{s,M}$ is the extensional stiffness of motors. F-actins and the center points of motors always exist two separate xy planes whose inter-distance is 13.5 nm. Note that a lipid bilayer is drawn in the schematic just to indicate that our motors are allowed move against a variable drag force, The bilayer is not part of our system. (B) Mean-square displacement (MSD) of motors measured as a function of the relative diffusion coefficient of motors ( $\bar{D}_M = D_M / D^*_M$). A higher MSD is indicative of more mobile motors. This confirms that imposing a higher diffusion coefficient to motors leads to more mobile motors.

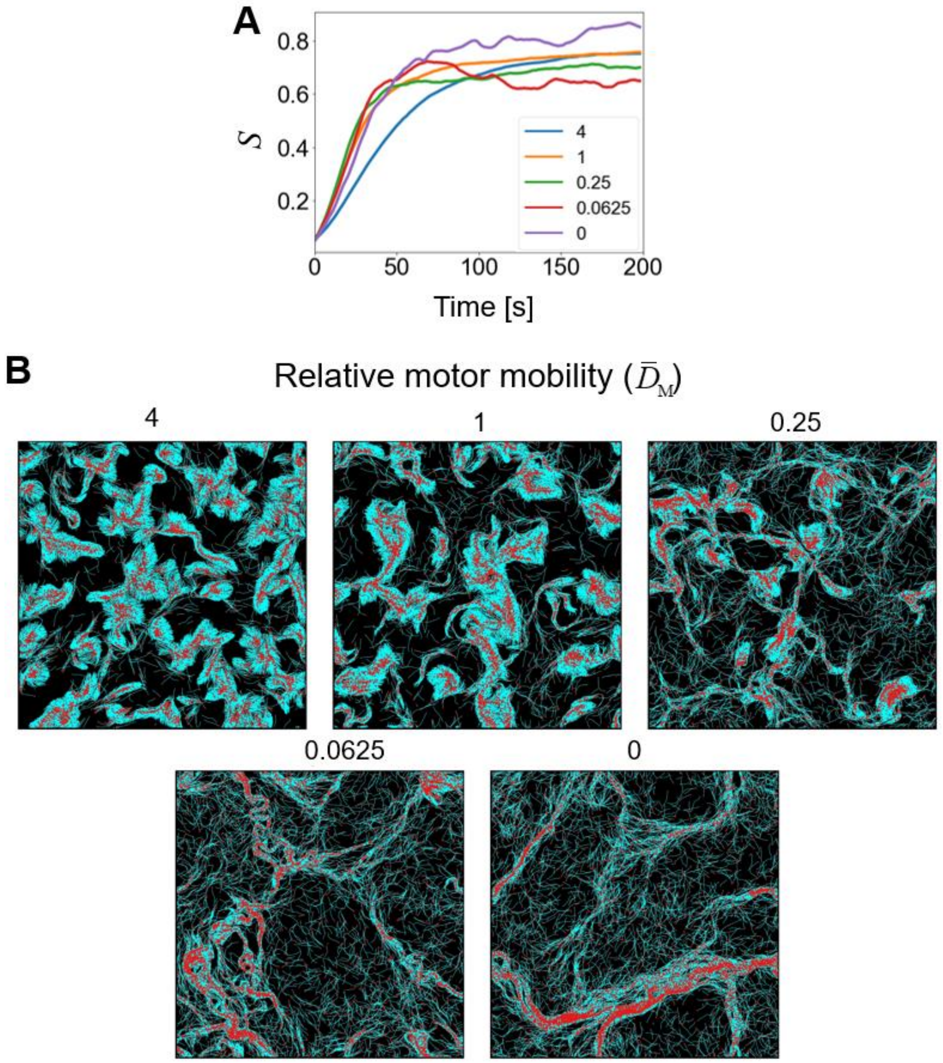


**Figure S2. Effects of motor mobility.** The motor mobility is defined by the relative diffusion coefficient of motors ($\bar{D}_M = D_M / D^*_M$). Numbers indicate the values of $\bar{D}_M$. (A) Nematic order parameter, $S$, depending on $\bar{D}_M$. There was no clear trend in $S$ unlike one observed in the polar order parameter, $P$. (B) Snapshots of F-actins (cyan) with their pointed end (red) labeled in simulations with different $\bar{D}_M$. In case of clusters, the pointed ends of F-actins tended to be localized near the centers of the clusters.

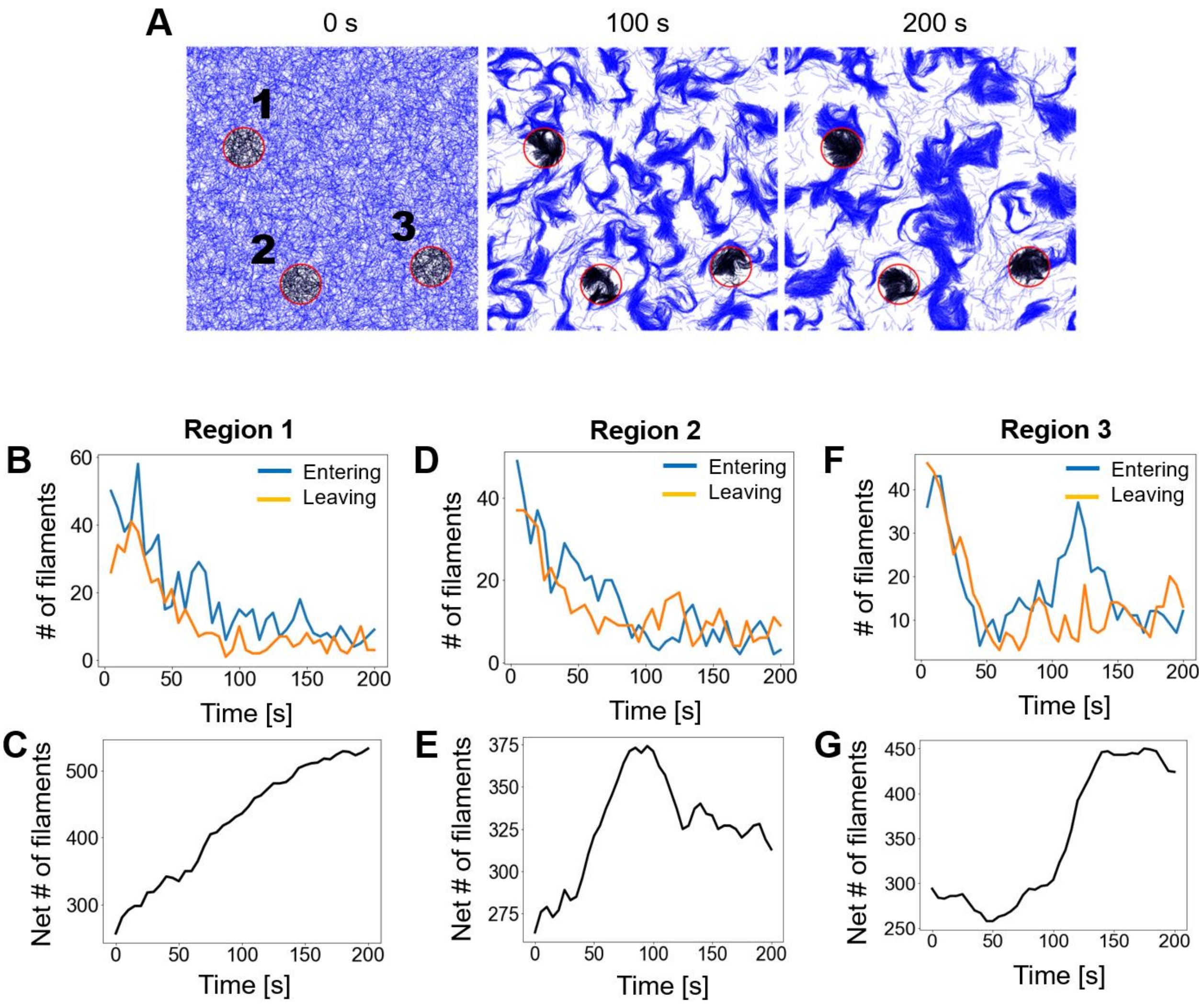


**Figure S3. Analysis of F-actins entering and leaving clusters.** (a) Snapshot of the case with $\bar{D}_{\mathrm{M}}$ = 1 taken at 0 s, 100 s, and 200 s with three clustering regions numbered and highlighted by red circles. All F-actins within the regions are labeled in black, whereas the rest of F-actins are shown in blue. (b, d, f) The number of F-actins entering (blue) or leaving (orange) each clustering region. (c, e, g) The accumulated number of F-actins within each clustering region.

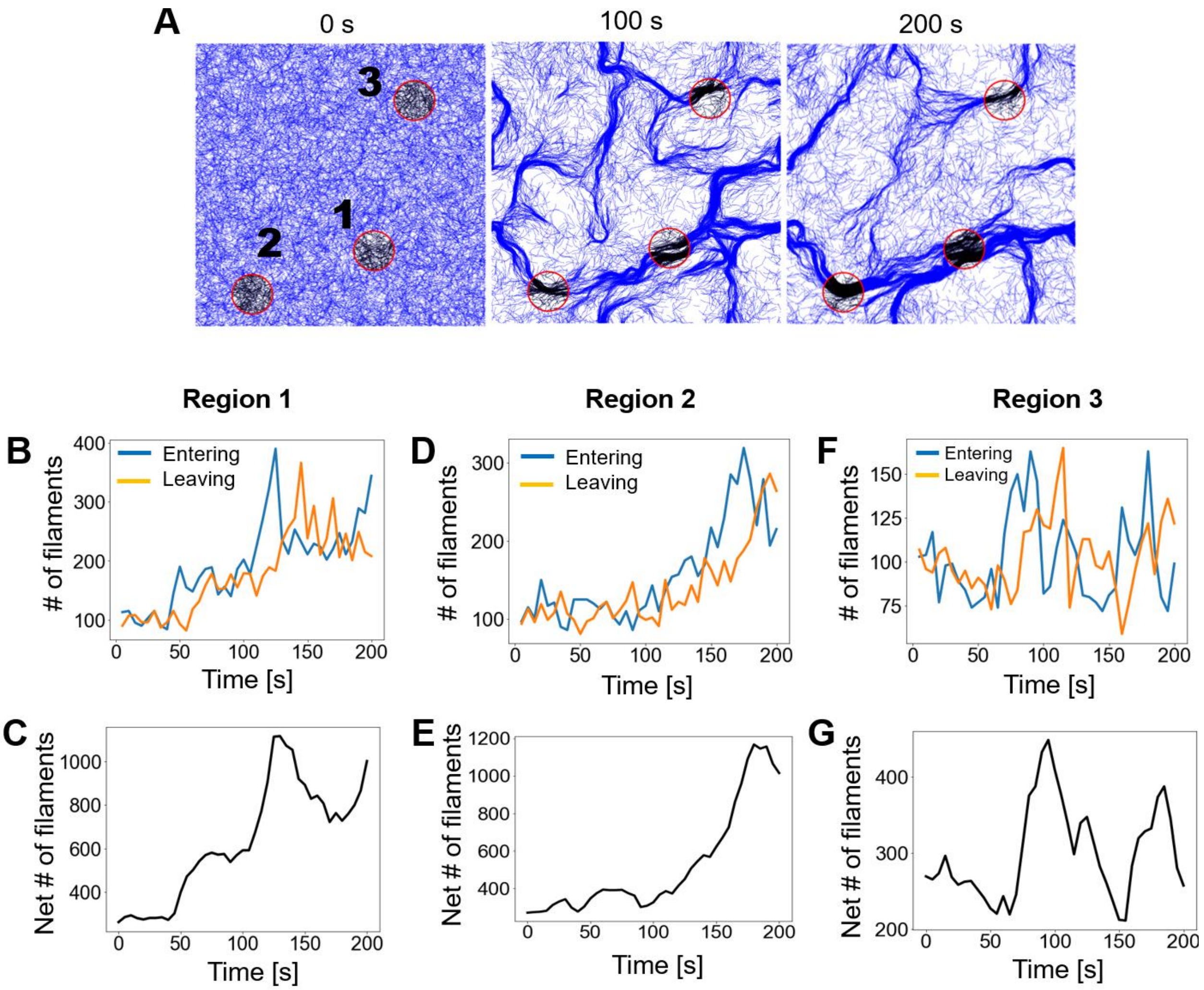


**Figure S4. Analysis of F-actins entering and leaving bundles.** (a) Snapshot of the case with $\bar{D}_{\mathrm{M}}$ = 1 taken at 0 s, 100 s, and 200 s with three bundling regions numbered and highlighted by red circles. All F-actins within the regions are labeled in black, whereas the rest of F-actins are shown in blue. (b, d, f) The number of F-actins entering (blue) or leaving (orange) each bundling region. (c, e, g) The accumulated number of F-actins within each bundling region.

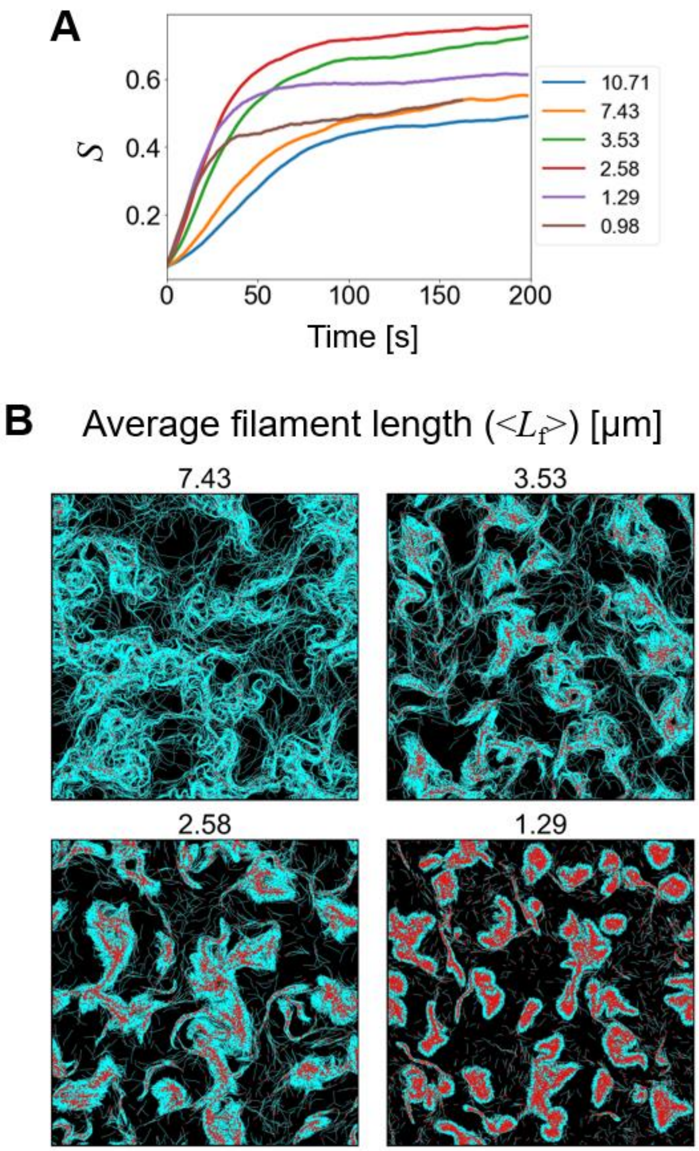


**Figure S5. Effects of average filament length (<$L_f$>).** Numbers indicate the values of <$L_f$> in µm. (A) Nematic order parameter, $S$, with different <$L_f$>. (B) Snapshots of F-actins (cyan) with their pointed end (red) labeled in simulations with different <$L_f$>. In case of small clusters formed by short F-actins, the localization of the pointed ends near the centers of the clusters is apparent.

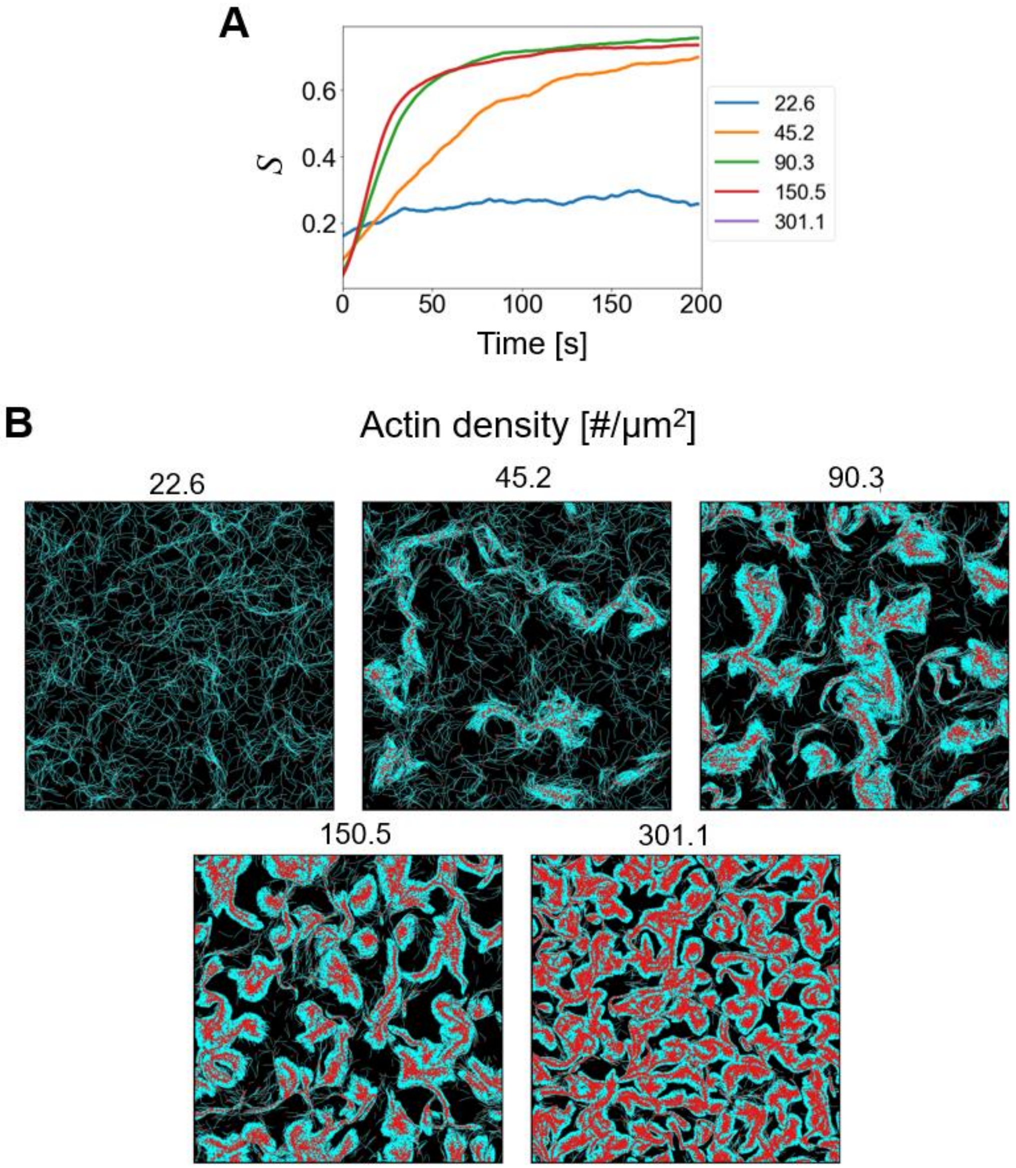


**Figure S6. Effects of actin density.** The actin density is defined by the number of actin monomers per μm$^2$. Numbers indicate the values of actin density. (A) Nematic order parameter, *S*, with different actin density. (B) Snapshots of F-actins (cyan) with their pointed end (red) labeled in simulations with different actin density. With higher actin density, the localization of the pointed ends near the center of structures is more apparent.

#### MOVIE CAPTIONS

**Movie S1.** Cases with different motor mobility defined by the relative diffusion coefficient of motors, $\bar{D}_{M} = D_{M} / D^{*}_{M}$. The numbers above five cases indicate the values of $\bar{D}_{M}$. The orientations of F-actins are visualized via the color scaling.

**Movie S2.** Magnified views of a clustering structure observed in the case with $\bar{D}_{M} = 1$ and a long bundle observed in the case with $\bar{D}_{M} = 0$. The orientations of F-actins are visualized via the color scaling.

**Movie S3.** Cases with different average F-actin length as indicated by numbers in the unit of μm. The orientations of F-actins are visualized via the color scaling.

**Movie S4.** Cases with different actin concentration as indicated by numbers in the unit of the number of actin monomer per $\mu m^2$. The orientations of F-actins are visualized via the color scaling.

### SUPPORTING TABLES

**Table S1. List of parameters employed in the model.** For some of the parameters, references are provided if the parameters were determined based on specific previous studies.

| Symbol | Definition | Value |
|---|---|---|
| $r_{0,A}$ | Length of an actin segment | $1.4\times10^{-7}$ [m] |
| $r_{c,A}$ | Diameter of an actin segment | $7.0\times10^{-9}$ [m] (2) |
| $\theta_{0,A}$ | Bending angle formed by adjacent actin segments | 0 [rad] |
| $\kappa_{s,A}$ | Extensional stiffness of F-actin | $1.69\times10^{-2}$ [N/m] |
| $\kappa_{b,A}$ | Bending stiffness of F-actin | $2.64\times10^{-19}$ [N·m] (3) |
| $\kappa_{r,A}$ | Strength of repulsive forces between F-actins | $3.38\times10^{-4}$ [N/m] |
| $r_{0,M}$ | Length of a motor segment | $1.35\times10^{-8}$ [m] |
| $r_{c,M}$ | Diameter of a motor segment | $1.0\times10^{-8}$ [m] |
| $\kappa_{s,M}$ | Extensional stiffness of a motor segment | $1.0\times10^{-3}$ [N/m] |
| $\Delta t$ | Time step | $1.15\times10^{-5}$ [s] |
| $\mu$ | Viscosity of a surrounding medium | $8.6\times10^{-1}$ [kg/m·s] |
| $k_B T$ | Thermal energy | $4.142\times10^{-21}$ [J] |
| $C_A$ | Actin concentration | 15 – 200 [μM] |
| $C_M$ | Motor concentration | 6 [μM] |
| $\langle L_f \rangle$ | Average length of F-actins | 1.29 – 7.43 [μm] |

**Table S2. List of the values of parameters used for "parallel cluster model." (4, 5)**

| Symbol | Definition | Value |
|---|---|---|
| $k_{01}$ | A rate from unbound to weakly bound state | 40 [$s^{-1}$] |
| $k_{10}$ | A rate from weakly bound to unbound state | 2 [$s^{-1}$] |
| $k_{12}$ | A rate from weakly bound to post-power-stroke state | 1000 [$s^{-1}$] |
| $k_{21}$ | A rate from post-power-stroke to weakly bound state | 1000 [$s^{-1}$] |
| $k_{20}$ | A rate from post-power-stroke to unbound state | 80 [$s^{-1}$] |
| $F_0$ | Constant for force dependence | $5.04\times10^{-12}$ [N] |
| $E_{pp}$ | Free energy bias toward the post-power-stroke state | $-60\times10^{-21}$ [J] |
| $E_{ext}$ | External energy contribution | 0 [J] |
| $d$ | Step size | $7\times10^{-9}$ [m] |
| $k_m$ | Spring constant of the neck linkers | $1.0\times10^{-3}$ [N/m] (= $\kappa_{s,M}$) |
| $N_h$ | Number of heads represented by a motor arm | 4 |
